# A dual-receptor mechanism between integrins and ACE2 widens SARS-CoV-2 tissue tropism

**DOI:** 10.1101/2022.01.02.474028

**Authors:** Danielle Nader, Timothy E. Gressett, Md Lokman Hossen, Prem Chapagain, Steven W. Kerrigan, Gregory Bix

**Author notes:** Correspondence to Prof’s Gregory Bix and Steve W. Kerrigan. In response to Zech et al. Nature Communications https://doi.org/10.1038/s41467-021-27180-0 (2022).

## Abstract

In addition to the ACE2 receptor, SARS-CoV-2 binds to integrins to gain host cell entry and trigger pro-inflammatory integrin-mediated signalling cascades. Integrins, therefore, are likely candidates for a dual-receptor mechanism with ACE2 to explain the increased infectivity seen in SARS-CoV-2 models. As integrins are primarily expressed in vasculature and persistent vasculopathy is seen in COVID-19, examining the role of endothelial integrin involvement is crucial in uncovering the pathophysiology of SARS-CoV-2.

---

The ability of SARS-CoV-2 to bind to host cells is a critical driver of infection and zoonotic transmission. Recently, Zech *et al*. demonstrated evidence which suggests that an amino acid mutation at position 403 (R403) of the SARS-CoV-2 spike protein is a critical factor for bat coronavirus spike protein to bind human ACE2^1^. However, we suggest the exigent importance of another major cell receptor, *integrins*, which also bind to R403 and which a growing body of literature supports their involvement in driving SARS-CoV-2 infection.

In their recent article, Zech and colleagues use computational analysis to study the electrostatic interactions between the SARS-CoV-2 spike protein trimer and ACE2 receptor and conclude that the R403 site is in close proximity with the E37 residue of ACE2. We believe, however, that this approach may have been assumptive. Iterative analysis reveals that the bond lengths between the speculated interacting side chains of these amino acids are 7.68Å and 8.1Å (Fig. 1A,B). We note that bond lengths greater than 4Å are considered to be energetically unfavourable, suggesting that this interaction is unlikely to form a strong bond *in vivo*^2^. Our group, in fact, has shown that bond lengths between the R403 site and integrin surface residues reveal close proximity and contain strong electrostatic bonds that are energetically favourable for *in vivo* interactions (Fig. 1C). Furthermore, a significant increase in binding affinity with residue D218 at 2.8Å occurs in the K403R mutation from the SARS-CoV predecessor, alongside Q180 at 3.7, and D150 at 3.3A^3^. It is also well documented that RGD motifs— found at spike protein R403, G404, D405— have exceedingly high affinity for integrins, which surpass any putative interaction with ACE2 yet reported to occur *in vivo*^4^. This suggests that when exposed to both ACE2 and integrins on a target cell, the solvent-exposed R403 would preferentially bind to integrins as part of an RGD-dependent interaction, rather than weakly interact with ACE2.

**Fig. 1.**
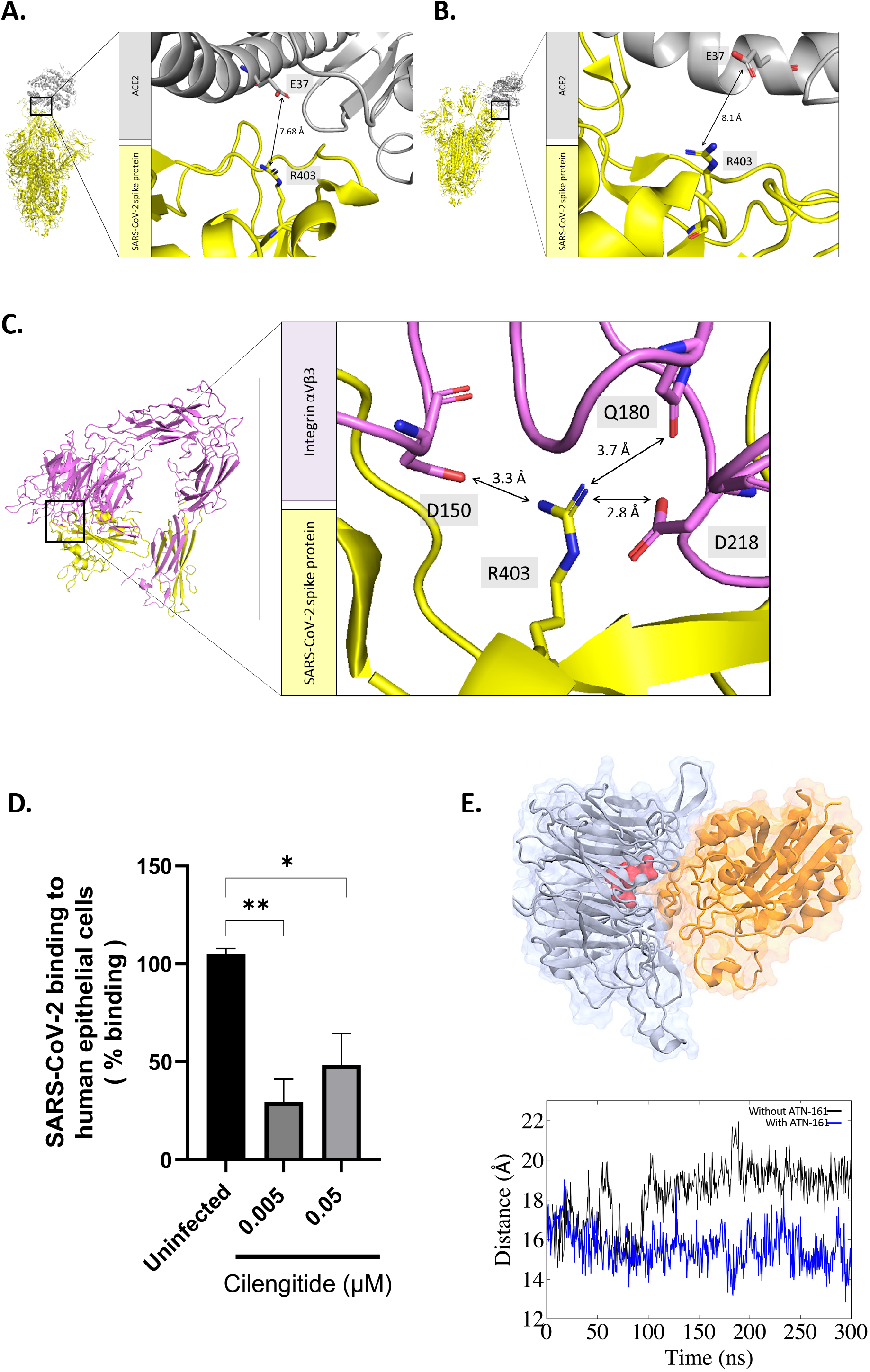
Integrins are a novel receptor of SARS-CoV-2 spike protein through the R403 mutation. **A.** Molecular modelling of human ACE2 in complex with SARS-CoV-2 spike protein (PDB: 7KNB) highlighting bond distance between residues E37 and R403, respectively. Protein structures were repaired and bond lengths were quantified using the PyMol bond measurement wizard then validated using the CCG MOE distance tool. **B.** Similarly, modelling of human ACE2 in complex with SARS-CoV-2 spike protein of the B.1.617.2. variant of concern (PDB: 7V8A). **C.** Docked integrin αVβ3 with receptor binding domain of SARS-CoV-2 spike protein, highlighting bond distance between integrin residues Q180, D150, and D218 with spike R403. **D.** Human epithelial Caco-2 cells were infected with SARS-CoV-2 for 24 hours as described previously3. Cells treated with Cilengitide displayed significantly reduced viral binding and infectivity (P<0.05). Data represents the mean of three independent experiments (±SEM). **E.** Docked integrin α5β1 with integrin antagonist ATN-161 (red). Center of mass distance between the RGD-binding residues D227 in α5 and S134 in β1 (right panel). The RGD-binding distance for the complex with ATN-161 shows a more restricted site compared to the complex without ATN-161.

Care must be taken, also, to utilize the proper cell types which may better model the influence of integrin binding alongside ACE2 when examining the role of R403 in SARS-CoV-2 adherence. Zech *et al*. used cells in their *in vitro* infection assay which did not express integrins and which we believe did not completely model the influence of integrin binding to R403. These cells include human epithelial colorectal Caco-2, embryonic kidney HEK293T, alveolar basal epithelial A549, and submucosal gland Calu-3. A first consideration which merits review is that Zech *et al*. did not use endothelial cells, which we believe would more accurately model the well-documented vascular dysregulation that occurs in COVID-19. In addition, the Caco-2 epithelial cells used lacked sufficient integrin heterodimer expression – we note the distinct absence of β3 RNA expression, as well as notably reduced levels of αV, α5, and β1^5^. These are integrin subunits which form heterodimeric pairs and have been demonstrated both *in silico* and *in vitro* to be key mediators in SARS-CoV-2 related vasculopathy and infection^3,6–10^. Thus, although the authors showed the enhanced effect that T403R had on viral attachment and entry, they did not, in our view, establish ACE2 as the determinant receptor involved in mediating this effect.

A second consideration was the 6-fold increase in viral replication in Caco-2 cells as compared to Calu-3 and HEK293T overexpressing ACE2. We note that the integrin subunit αV is expressed in Caco-2 cells, albeit moderately, and our molecular modelling has revealed several contact points which exist between spike protein R403 and αV at energetically favourable distances for *in vivo* interaction (Fig. 1C). Additionally, there was a 2.5-fold increase in viral replication in A549 cells, which express the integrin subunit β1 –significant, again, as recent *in silico* candidate screening has firmly identified β1 as putative receptors across these epithelial cells^11^. Therefore, we believe that integrins are likely contributors to the increase in SARS-CoV-2 binding to Caco-2, and to some extent A549 cells.

Lastly, although the integrin inhibitor ATN-161 is known to bind integrin αVβ3, it primarily targets α5β1. This could elucidate why ATN-161 was deemed ineffective when used in SARS-CoV-2 infected Caco-2 cells, as they do not reliably express the α5β1 heterodimer. Caco-2 cells do moderately express integrin subunit αV, however, and our group has shown that Cilengitide, an RGD cyclic pentapeptide which targets αV, significantly reduces SARS-CoV-2 attachment and epithelial infectivity *in vitro* using Caco-2 (Fig. 1D). Additionally, our molecular modelling predicts that R403 heavily interacts with residues of αV rather than β3, such that Cilengitide likely retains effectiveness in the absence of a whole integrin heterodimer. Likewise, molecular dynamics simulations of ATN-161 binding with α5β1 show that it binds the inner cavity of α5 and can alter the RGD binding site, reducing the availability of spike R403 to bind to this integrin (Fig. 1E). Therefore, ATN-161 would have been more appropriate to use on cells that reliably express α5β1 integrins. Cilengitide, similarly, would have been better suited to interrogate the role of integrins in viral infectivity in Caco-2 cells. In fact, our group was the first to publish evidence that supports both ATN-161 and Cilengitide inhibit SARS-CoV-2 binding and infectivity in the vasculature, where integrins are widely expressed and function to control cellular permeability and inflammation^3,6^. More importantly, we were also the first to demonstrate that inhibition of integrin α5β1 and αvβ3 *in vivo* can reduce both SARS-CoV-2 viral load and pathological complications.^12^

Recent data has identified binding sites within the intracellular tails of both integrins and ACE2 that serve to promote receptor interplay and cooperative signalling^13,14^. Thus, intracellular signalling dynamics may also clarify how integrins and ACE2 act as co-receptors for SARS-CoV-2 spike protein. Structural measurements and molecular modelling of these intracellular structures bound to their respective ligands reveal that integrins and ACE2 exist relatively close in space (Fig 2 A,B). Therefore, the highly flexible spike protein trimer on a single virion could interact with both integrins and ACE2 simultaneously. This is striking as it suggests the occurrence of a dual-receptor mechanism which may elucidate how R403 interacts with both integrins and ACE2 to increase SARS-CoV-2 virulence.

**Fig. 2.**
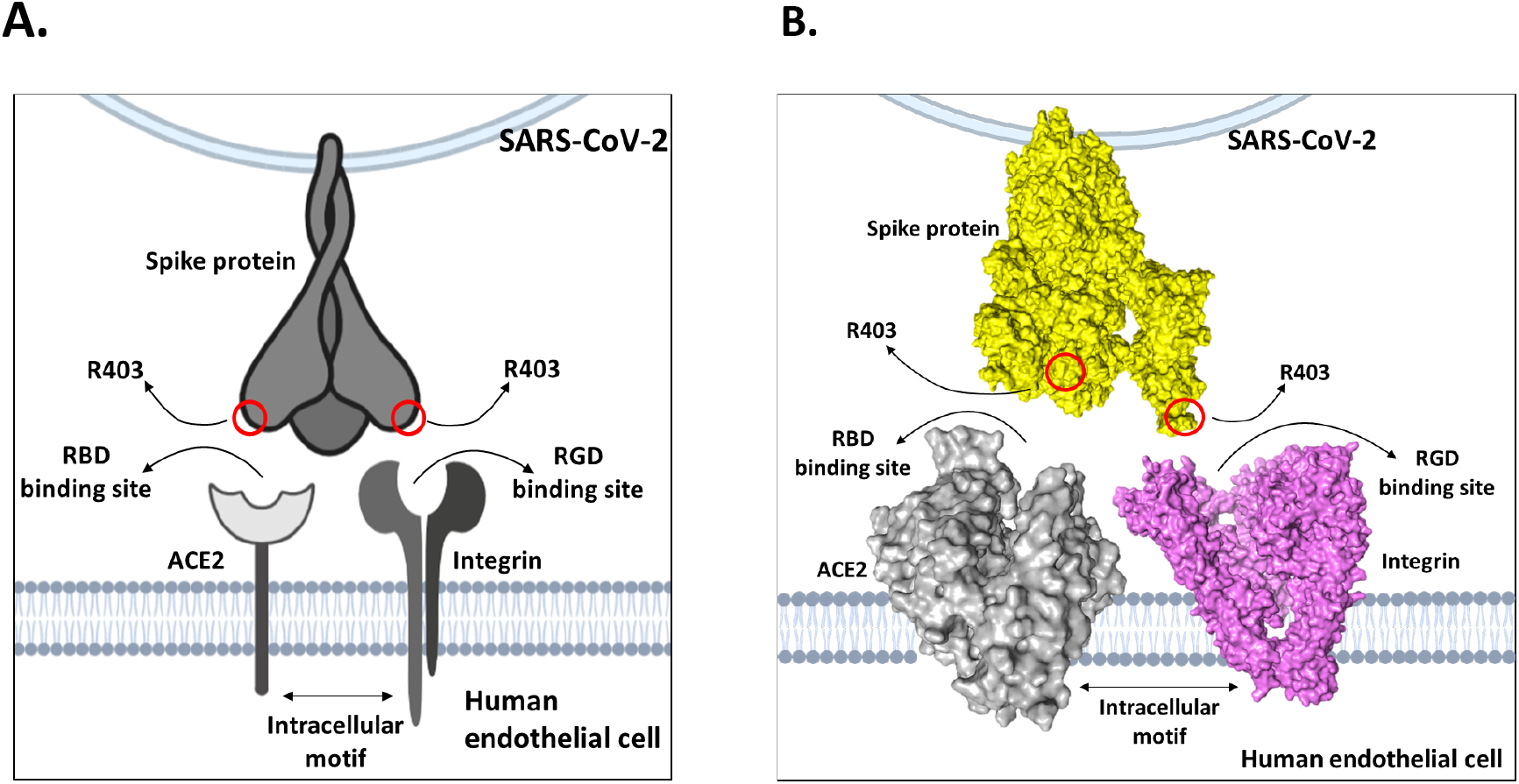
Model of dual-receptor mechanism between integrins and ACE2 to mediate enhanced tissue tropism of SARS-CoV-2 spike protein **A.** Schematic diagram illustrating SARS-CoV-2 spike protein trimer present on a single virion, in contact with a human endothelial cell. Each trimer contains a receptor binding domain carrying a surface-exposed R403 residue, highlighted in red. Arrows indicate locations of RGD and RBD-binding sites on integrin and ACE2, respectively. Interacting intracellular motifs are present within the cystolic integrin and ACE2 tails, placing them in close proximity. **B.** Surface maps of SARS-CoV-2 spike protein, integrin αVβ3, and ACE2 created in PyMol. High flexibility of the spike protein trimer may allow interaction with both receptors simultaneously due to the close nature of the two receptors.

Broadly, whilst we accept that R403 may support spike protein attachment to ACE2 in epithelial cells, we believe that the critical evolutionary benefit of this mutation may be better explained by its potential interaction with integrins. Our modelling indicates that it is unlikely that a strong electrostatic interaction occurs between spike protein R403 and ACE2 receptor E37 *in vivo*. The energetically significant contacts found between R403 and integrins, however, is compelling. Both *in vivo* and *in vitro* evidence published by ourselves and others supports integrins as novel receptors of SARS-CoV-2, making integrin antagonists attractive potential therapeutic interventions to reduce viral adherence and infection^3,6–10,12^. Finally, it has been reliably observed that COVID-19 causes severe endothelial injury^15^. It is therefore critical to investigate the role R403 plays in SARS-CoV-2 dissemination, and continue research efforts using assays which account for the unique genomic expression profiles inherent to the vasculature that include integrin expression. Continued *in vivo* work examining the ability of ATN-161 and Cilengitide as novel therapeutic interventions for SARS-CoV-2 is also urgently warranted.

We propose that R403 has evolved to hijack endothelial integrins to drive intracellular signalling cascades that mediate cellular permeability and vascular damage in COVID-19, and to some extent, binding ACE2 in the epithelium. We agree with Zech *et al*. that the R403 amino acid substitution on the spike protein has played a critical role in mediating the pandemic spread. We propose, however, that a dual-receptor mechanism between integrins and ACE2 ultimately widens SARS-CoV-2 tissue tropism (Fig 2 A,B), and therefore encourages significantly more efficient transmission as seen epidemiologically.

## Methods

### Protein structural analysis

Human ACE2 receptor in complex with SARS-CoV-2 spike protein (PDB: 7KNB) and B.1.617.2. variant of concern (PDB: 7V8A) were downloaded from RCSB. Structures were energetically repaired in Yasara using the FoldX suite and visually compared using PyMol and CCG Molecular Operating Environment (MOE). Distances between the structural coordinates of the spike protein RGD motif (403-405) and ACE2 E37 residue were quantified using the PyMol bond measurement wizard and validated with the MOE distance tool.

### Molecular docking and Molecular Dynamics (MD) simulations

PHSCN peptide was docked to the open conformation of human integrin α5β1 (PDB ID: 7NWL)^16^ using HPEPDOCK server^17^ as well as AutoDock Vina^18^. Both tools yielded similar binding poses in the β-propeller domain. Only the best ranked binding pose obtained from AutoDock Vina was used for MD simulations. Two systems of α5’s β-propeller domain were prepared for MD simulation, one with PHSCN and one without, using solution builder interface of CHARMM-GUI^19^ with CHARMM36 force field. Both systems were simulated at 300K using GPU version of NAMD 2.14^20,21^. A 10,000 steps minimization and 250 ps equilibration were performed. Finally, a 300 ns unconstrained production run was performed with 2 fs/step for each system. The visualization and data analysis were performed using VMD 1.9.3^22^. Integrin αVβ3 (PDB: 1L5G) docked with receptor binding domain of SARS-CoV-2 spike protein (PDB: 6M0J) was performed in PyMol as described previously^18^.

### In vitro assessment of Cilengitide inhibition of SARS-CoV-2 binding

An *in vitro* para-nitrophenyl phosphate binding assay was performed using human colorectal adenocarcinoma Caco-2 cells and formaldehyde-fixed SARS-CoV-2 to investigate the ability of RGD cyclic pentapeptide compound Cilengitide (0.05 and 0.005 μM) to reduce viral adherence, as described previously^18^. Briefly, Caco-2 cells were infected with Human 2019-nCoV strain 2019-nCoV/Italy-INMI1 at a Multiplicity of Infection of 0.4. Following two hours, a para-nitrophenyl phosphate solution in lysis buffer was added to infected cells, where the fluorescent signal was measured at 405 nm to quantify cell-viral binding (%). SARS-CoV-2 samples were obtained from the European Virus Archive Global (Ref. no: 008V-03893) and experiments were carried out under Biosafety level 2 guidelines.

### Statistical analyses

Data is represented as mean ± standard error of the mean. Experiments were carried out in triplicate with a minimum of three independent experiments. Statistical differences between groups were assessed using one-way ANOVA followed by Tukey’s multiple comparison post-hoc test. P-value < 0.05 was considered to be significant, indicated with asterisks: *P<0.05, **P<0.01.

## Acknowledgements

We would like to acknowledge the following individuals for their support and useful conversations in putting together this manuscript: Tione Buranda, Juan Pablo Robles, and Amruta Narayanappa.

## Contributions

D.N. and T.G. wrote and edited the manuscript. G.B. and S.K. conceived the manuscript and provided critical review. Figure 1 D was created by P.C. and M.L.H. The remaining Figures 1 and 2 were created by D.N. with inputs from all authors. All authors equally reviewed and edited final manuscript.

## Ethics declarations

Competing interests

The authors declare no competing interests.

